# Modelling and analysis of bacterial tracks suggest an active reorientation mechanism in *Rhodobacter sphaeroides*

**DOI:** 10.1101/001917

**Authors:** Gabriel Rosser, Ruth E. Baker, Judith P. Armitage, Alexander G. Fletcher

## Abstract

Most free-swimming bacteria move in approximately straight lines, interspersed with random reorientation phases. A key open question concerns varying mechanisms by which reorientation occurs. We combine mathematical modelling with analysis of a large tracking dataset to study the poorly-understood reorientation mechanism in the monoflagellate species *Rhodobacter sphaeroides.* The flagellum on this species rotates counterclockwise to propel the bacterium, periodically ceasing rotation to enable reorientation. When rotation restarts the cell body usually points in a new direction. It has been assumed that the new direction is simply the result of Brownian rotation. We consider three variants of a self-propelled particle model of bacterial motility. The first considers rotational diffusion only, corresponding to a non-chemotactic mutant strain. A further two models also include stochastic reorientations, describing ‘run-and-tumble’ motility. We derive expressions for key summary statistics and simulate each model using a stochastic computational algorithm. We also discuss the effect of cell geometry on rotational diffusion. Working with a previously published tracking dataset, we compare predictions of the models with data on individual stopping events in *R. sphaeroides*. This provides strong evidence that this species undergoes some form of active reorientation rather than simple reorientation by Brownian rotation.

## 1 Introduction

The motile behaviour of bacteria underlies many important aspects of their actions, including pathogenicity, foraging efficiency and biofilm formation. The study of bacterial motility is therefore of biomedical and industrial importance, with implications in the control of disease [1] and biofouling [2]. Bacteria swim by rotating semi-rigid helical flagella, powered by the movement of ions through a transmembrane rotary motor. The motor can switch between clockwise and counterclockwise rotation, the change in torque transforming the wavelength and handedness of the filament helix. Owing to their small size, a flagellate bacterium’s free-swimming behaviour is characterised by a Reynolds number on the order of 10^−5^ [3]. Approximating a bacterium by a 1 *μ*m-diameter sphere moving at 20 *μ*m s^−1^, the coasting distance upon instantaneous cessation of the flagellar motor is around 4 × 10^−12^m, or 4 millionths of the bacterium’s diameter [4]. This calculation demonstrates that, at such low Reynolds numbers, viscous effects dominate over inertia, and objects are brought to a halt almost immediately.

Despite having extremely small coasting distances, bacteria are never completely stationary as they exhibit Brownian motion [3]. This motion arises from the net force acting on a bacterium as a result of the large number of collisions with surrounding molecules in the liquid. Although the time-averaged displacement as a result of these collisions is zero, the instantaneous net force is a random property, causing the bacterium to exhibit ceaseless small movements. Such Brownian buffeting leads to two related processes: translational and rotational diffusion (see Figure 2). Translational diffusion leads to a shift in the bacterium’s centre of mass over time, whereas rotational diffusion leads to a reorientation of the bacterium about its centre of mass.

**Figure 1.**
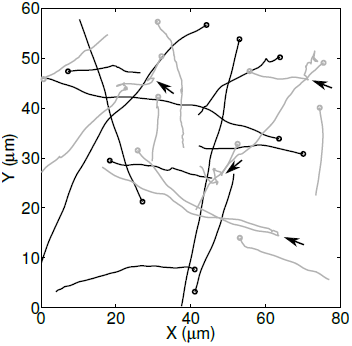
Tracks from the experimental dataset used in this study. Black lines correspond to the non-chemotactic mutant, grey lines show wildtype tracks. Several reorientation events are indicated by black arrows.

**Figure 2.**
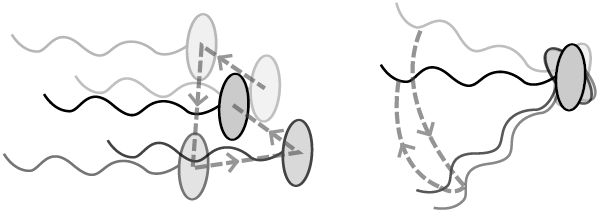
Illustration of a single bacterium undergoing translational (left) and rotational (right) diffusion. Increasing transparency represents position and orientation in the more distant past. Dashed lines trace the trajectory of the cell centroid (left) or a point on the flagellum to show the angle changes (right).

A further consequence of a bacterium’s microscopic size is its inability to detect concentration changes in nutrients or toxins over the length of the cell body. This limitation affects motile behaviour. Many species, such as the multiflagellate *Escherichia coli*, swim in a series of approximately straight ‘runs’ interspersed by reorientating ‘tumbles’. During a run, the flagellar motors turn counter-clockwise, causing the helical flagella to form a rotating bundle that propels the bacterium forward. Tumbles occur when one or more motors reverse their rotation, disrupting the flagellar bundle and causing the cell to reorient in a random manner [5]. A related mechanism exists in the uniflagellate bacterium *Rhodobacter sphaeroides*, in which reorientations are, instead, caused by stopping the flagellar motor [6]. Upon ceasing to rotate, the flagellum undergoes a conformational change, transforming from a functional semi-rigid helix to a short wave length, high amplitude coil against the cell body [7]. This leads to reorientation by a mechanism that is not yet well understood.

By modulating the frequency of reorientation events based on a temporal comparison of the concentration of various chemicals in their immediate surroundings, bacterial species such as *E. coli* and *R. sphaeroides* may bias their movements towards sources of nutrients or other substances (collectively termed chemoattractants), in a process called chemotaxis [8]. The biochemical pathways responsible for chemotaxis in *R. sphaeroides* are less fully elucidated than those in *E. coli*, and are known to be more complex [9]. In combination with experimentation, mathematical modelling can help gain insight into patterns of bacterial motility and chemotaxis.

A common approach to modelling run-and-tumble motility involves a velocity jump (VJ) process, in which the effect of rotational diffusion is neglected and runs are assumed to occur in straight lines, interspersed with stochastic reorientation events [10]. Each running phase occurs with a constant velocity drawn from an underlying distribution. This model has been extended to include key intracellular signalling pathways in order to study the response to changes in chemoattractants in the surrounding environment [11].

While the VJ approach provides a reasonable first approximation of run-and-tumble motility, in reality rotational Brownian buffeting prevents bacteria from swimming in perfectly straight runs. In order to bias their movements towards nutrient sources, bacteria must swim sufficiently rapidly to detect a change in nutrient concentration before rotational diffusion randomises the direction of swimming. Purcell [4] has described this requirement as the need to ‘out-swim diffusion’.

An important early discussion of the effect of rotational diffusion on bacterial motility was given by Berg [3], who modelled a bacterium as an ellipsoid in order to present approximate expressions for the variation of the strength of rotational diffusion with cell dimensions. The author derived an approximate expression for the effective diffusion coefficient of a mutant bacterium that swims at a constant speed without reorientating.

In other work, Mitchell [12] used a self-propelled particle model to analyse the effect of cell size and swimming speed on the efficiency and feasibility of bacterial motility in marine environments. This work suggested that the extent of rotational diffusion is independent of a bacterium’s swimming speed, but varies as *r*^−3^ with its effective radius, *r*. An important conclusion of this work is that smaller cells must swim more rapidly in order to undergo chemotaxis. In a further study, Mitchell [13] found that incorporating frictional effects of the flagellum into his model diminished the extent of rotational diffusion, as this increases the viscous drag acting on the bacterium, termed ‘flagellar stabilization’. Dusenbery [14] carried out a similar analysis to Mitchell [13], predicting that planktonic organisms with radii below around 0.6*μ*m are unlikely to gain any advantage from locomotion and verifying this prediction through a systematic investigation of known bacteria [14]. It is, however, unclear whether the more general prediction of Mitchell [13], that swimming speed should correlate with cell size, holds in nature; a diverse study of marine bacteria carried out by Johansen et al. [15] failed to find any such correlation.

An important open question in this field concerns the varying mechanisms by which bacteria reorientate. Discussing the implications of their model of bacterial motility, Mitchell and Kogure [16] hypothesise that *R. sphaeroides* is too large to reorientate efficiently by rotational Brownian diffusion alone. Further experimental evidence for an active reorientation mechanism in *R. sphaeroides* was obtained by Armitage et al. [7] using differential interference contrast microscopy. However, more recent bead assays in *R. sphaeroides* have provided contradictory evidence, suggesting that the motor does indeed stop [6]. These conflicting results are likely to be due in part to significant differences in experimental protocols. Free-swimming methods assess the behaviour of flagella rotating with low loads on the filament, while experiments on tethered cells, in which a bead is attached to the flagellum, may artificially increase the load on the filament, leading to different results. Indeed recent analysis suggests that the number of stators engaged in actively driving rotation increases with load, and long term increased loads on the filament may affect observation of reorientation events, which are usually very short-lived [17].

To date, a major obstacle in making further progress in this area is the difficulty in obtaining quantitative data on the free-swimming behaviour of *R. sphaeroides* under normal conditions. A common approach for obtaining quantitative data on bacterial motility involves the tracking of free-swimming cells using video microscopy. However, the analysis of such data remains challenging, in particular the identification of reorientation events in tracks in the face of various sources of noise within the data. A novel automated, non-parametric method of analysing large bacterial tracking datasets was recently proposed by Rosser et al. [18], who demonstrated its validity and reliability through computational simulations. Using video microscopy, the authors recorded and analysed thousands of cell tracks for *R. sphaeroides* and *E. coli*. They found that *R. sphaeroides* exhibits directional persistence over the course of a reorientation event. Such tracking data represent an excellent opportunity to investigate the role of Brownian rotational buffeting in bacterial motility, in addition to permitting a novel approach to the study of the poorly-understood reorientation mechanism in *R. sphaeroides*. In particular, the study provides rich data from a non-chemotactic mutant strain that is unable to reorientate, permitting a hitherto infeasible direct analysis of the role of rotational diffusion on a swimming *R. sphaeroides* bacterium. Furthermore, Rosser et al. [18] generated a large tracking dataset of wildtype *R. sphaeroides*, annotated to show where reorientation events occurred. To our knowledge, no previous studies have used bacterial tracking to investigate this phenomenon. A sample of the tracks in the tracking datasets is plotted in Figure 1.

The above work provides a foundation for the present study, in which we demonstrate a novel, phenomenological approach to study the reorientation mechanism in *R. sphaeroides* that exploits the availability of these tracking data. Our primary motivations for this study are twofold. First, we address the discrepancy between the idealised VJ description of bacterial motility and the non-straight line runs evident in experimental tracks, such as those in Figure 1. Second, we wish to use the newly available tracking data as an alternative method for comparison with tethered cell studies to address the poorly understood phenomenon of reorientation in *R. sphaeroides*.

We use mathematical modelling, combined with novel analysis of the tracking dataset, to investigate the effect of rotational diffusion on bacterial motility. We consider three models of bacterial motility to analyse the role of rotational diffusion in the free-swimming behaviour of *R. sphaeroides*, and other monoflagellates with similar modes of motility. The first is a model developed by ten Hagen et al. [19] to describe the motion of a self-propelled particle that is subject to rotational diffusion only, representing the motion of a non-chemotactic mutant. The other models are novel and couple the self-propelled particle model with a simple VJ model of run-and-tumble motility.

In what follows, we describe the governing equations of motion. For each model we develop a stochastic simulation algorithm and derive expressions for key summary statistics: the first two moments of position and the mean squared angle change (MSAC). We next use the data generated by Rosser et al. [18] to estimate the translational and rotational diffusion coefficients of the observed bacteria and compare these with theoretical values. The effect of cell geometry on rotational diffusion is discussed using theoretical results; in particular we extend the work of Berg [3] by showing explicitly the dependence of the rotational diffusion coefficient on the dimensions of the ellipsoidal cell body. This enables a comparison with the result measured experimentally. Finally, we reconcile the phenomenon of rotational diffusion with the VJ process by describing the reorientation of a bacterium during a stopping phase in terms of rotational diffusion. By comparing each model with data on individual stopping events in *R. sphaeroides* obtained using our analysis method, we are thus able to show that *R. sphaeroides* needs more than Brownian rotational motion to reorient.

## 2 Materials and methods

Our mathematical models are based on the overdamped Langevin description of a bacterium as a self-propelled particle. The Langevin equation is a stochastic differential equation (SDE) that is commonly used to describe a particle undergoing Brownian motion in a liquid [20].

As our tracking data are captured in a two-dimensional (2D) focal plane far away from walls [18], our models assume bacterial motion to be confined to an infinite planar domain. In the case of an active propulsive force, we assume for simplicity that the bacterium is propelled forward at a constant speed of *c* = 40 *μ*m s^−1^, which is the approximate modal speed exhibited by wildtype *R. sphaeroides* [18]. Following ten Hagen et al. [19], we include only rotational diffusion in our models, as translational diffusion has negligible effect on the motion of a self-propelled particle when the rate of propulsion is sufficiently high, as is the case here. We consider the validity of these assumptions further in the *Discussion* section. Nonetheless, the magnitude of translational diffusion in non-motile bacteria provides a useful consistency check between experiment and theory, hence we also present relevant theoretical results on translational diffusion.

### 2.1 Bacterial cell geometry

As illustrated in Figure 3, the bacterium is modelled as either a sphere or an ellipsoid [21, 22] that is propelled by a single flagellum, represented by a rigid helix. In the case of a prolate ellipsoid, we denote the lengths of the axial and equatorial semi-principal axes by *a* and *b*, respectively (noting that the two equatorial semi-principal axes are degenerate), and quantify the geometry by the axial ratio, *ρ* = *a*/*b*, whose value is greater than one. When the cell body is ellipsoidal, the flagellum is attached at the midpoint of the long axis as is generally observed experimentally [23]. The unit vector in the axis that runs through the centre of the flagellum and the centre of the cell is the orientation vector ***μ***, whose angle to the horizontal is denoted *ϕ*. Since we consider only two-dimensional motion, we assume that ***μ*** lies in the (*x*, *y*) plane.

Note, however, that the cell body is three-dimensional in our models.

The translational and rotational diffusion coefficients of a general ellipsoid may be calculated using multiplicative adjustments to the translational and rotational frictional drag coefficients of a sphere of equivalent volume, respectively. These corrections are known as Perrin friction factors [24]. Berg [3] gives approximate expressions for these, however we use the exact forms here to ensure greater accuracy when *ρ* ≈ 1. Since we consider a bacterium with a medially attached flagellum (see Figure 3), we are concerned with rotation about the *equatorial* semi-principal axis; rotation about the axial axis would cause the bacterium to swim out of the plane, breaking the assumptions of the model. For a prolate ellipsoid, a valid expression for Perrin’s friction factor for rotation about the equatorial semi-principal axis is given by [24]

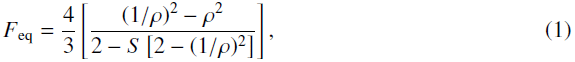

where for brevity we have defined a new variable

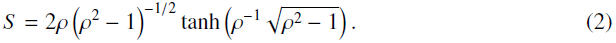

The volume of a prolate ellipsoid is given by 4*πab*^2^/3. To compute the rotational frictional drag coefficient for an ellipsoidal cell, we multiply the value for a sphere of equivalent volume by *F*_eq_.

**Figure 3.**
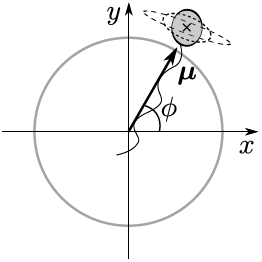
Schematic of a three-dimensional bacterium, confined to travel in the (*x*, *y*) plane, which undergoes rotational diffusion. Dashed lines represent alternative shapes for the cell body. The black cross indicates the centre of mass of the cell, which is calculated neglecting the effect of the flagellum.

### 2.2 Translational and rotational Langevin equations

The Langevin equation describing the translational diffusion of a particle in 2D is given by [20]

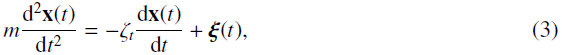

where **x**(*t*) = (*x*(*t*), *y*(*t*)) denotes the time-dependent position of the particle, *m* is its mass, *ζ*_*t*_ is the translational frictional drag coefficient and ***ξ***(*t*) = (*ξ*_*x*_(*t*), *ξ*_*y*_(*t*) is a random fluctuating force with independent components. The right-hand side of (3) represents the force exerted on the particle, which is equal to the sum of a deterministic viscous drag term and a force arising from the random collisions of the surrounding molecules in the liquid. The latter, ***ξ***(*t*), is given by a Gaussian white noise, with correlation function

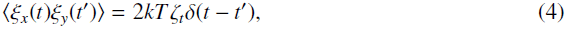

where *k* is Boltzmann’s constant, *T* is the temperature in units of kelvin, 〈·〉 denotes an ensemble average and *δ*(·) is the Dirac delta function.

The delta correlation summarised in (4) reflects the assumed separation of the viscous relaxation timescale and the timescale of the motion of the surrounding molecules. To justify this simplification, we first note that the motion of the molecules surrounding the particle are correlated on a timescale of 10^−12^ s [20]. We next calculate the approximate viscous relaxation timescale, given by *m*/*ζ*_*t*_ for a sphere of radius *r* = 1 *μ*m (the approximate diameter of many bacteria, including *R. sphaeroides* [25]) moving through buffer at room temperature. We may calculate a theoretical value of *ζ*_*t*_ for a sphere of radius *r* surrounded by a fluid with viscosity *η* using Stokes’ law,

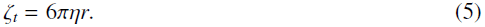

The viscosity of water at room temperature is approximately 10^−3^ Pa s, which gives *ζ*_*t*_ ≈ 10^−8^ kg s^−1^ for a cell in water. Assuming that the density of a cell is approximately equal to that of water at room temperature, *m* ≈ 10^−15^ kg. Hence the characteristic viscous relaxation timescale, *m*/*ζ*_*t*_, is around 10^−7^ s, several orders of magnitude slower than the correlation timescale for the motion of surrounding molecules. The white noise force is therefore a suitable approximation to the true fluctuating force acting on a Brownian particle. For comparison, the timescales on which we observe bacterial motility are on the order of 10^−3^ s or greater. Reinterpreting this derivation, we see that viscous forces dominate over inertia in the case of a bacterium subject to Brownian motion. We therefore approximate (3) by the overdamped translational Langevin equation

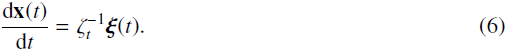

In the case of rotational diffusion, we consider an ellipsoidal body of uniform density with mass *m* and an orientation vector ***μ***(*t*), as described above and shown in Figure 3. The Langevin equations describing the rotational diffusion of this body in three dimensions are given by [20]

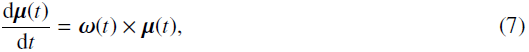

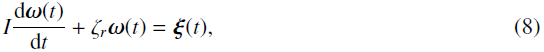

where *ζ*_*r*_ is the rotational drag coefficient, *I* = 2*mr*^2^/5 is the moment of inertia of the sphere, *ω* is the angular momentum and ***ξ***(*t*) is a white noise force as described above. The rotational drag coefficient for a sphere of radius *r* is given by the rotational analogue of 5,

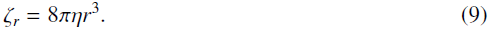

As before, inertial terms are small compared with viscous terms, so that *I* ≪ *ζ*_*r*_. We therefore approximate (7)–(8) by the overdamped rotational Langevin equation

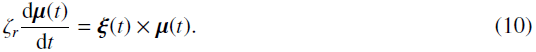

Finally, to reflect the assumed planar motion of the bacterium, we derive the simplified 2D form of equation (10) used by ten Hagen et al. [19]. In this case, we write the orientation vector as ***μ***(*t*) = (cos *ϕ*, sin *ϕ*, 0)^⊤^. Taking the dot product of equation (10) with the unit vector in the *x*-direction, we obtain

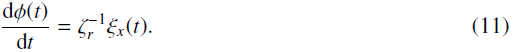

Here, and in the remainder of this work, we assume without loss of generality that *x*(0) = *y*(0) = *ϕ*(0) = 0, since the initial location and direction of travel of a bacterium are arbitrary.

### 2.3 Run-only model

In the run-only model, we assume that the propulsive force is active at all times, hence the bacterium travels at constant speed *c*. This is a simplified representation of the *R. sphaeroides* non-chemotactic mutant strain, which is unable to stop [6]. The governing overdamped Langevin equations for the run-only model are thus given by

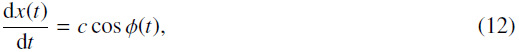

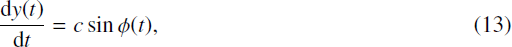

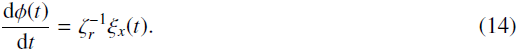

### 2.4 Run-and-stop model

In the run-and-stop model, we assume that the propulsion force undergoes stochastic switching events between running and stopping states as a Poisson process. Rotational diffusion continues to act regardless of the state of the propulsion force, hence the bacterium undergoes passive diffusional reorientation in the stopping phase. This model attempts to capture the details of a wildtype *R. sphaeroides* cell, assuming that stops occur by rotational diffusion alone.

To incorporate the effect of this random switching in the equations of motion of the self-propelled particle, we introduce a continuous time Markov process, *F*(*t*), on the state space {0,1}. The bacterium is in a stopping phase when *F*(*t*) = 0, and in a running phase when *F*(*t*) = 1. We denote the rate of switching from a run to a stop by *λ*_*s*_, and that of switching from a stop to a run by *λ*_*r*_. We assume that the stochastic processes *F*(*t*) and *ξ*_*x*_(*t*) are independent, and that the bacterium is initially running, so that *F*(0) = 1. The Langevin equations governing the motion of the running and stopping particle are thus given by

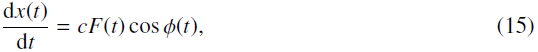

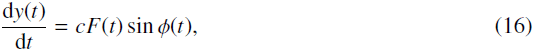

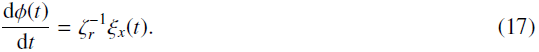

### 2.5 Run-and-active-stop model

In the run-and-active-stop model, we incorporate an additional stochastic rotational force of magnitude *α*, which acts as a multiplier to the rotational diffusion coefficient in stopping phases only, so that reorientation occurs more rapidly. The governing equations are thus given by

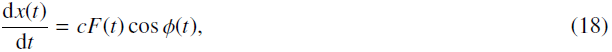

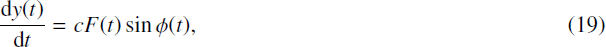

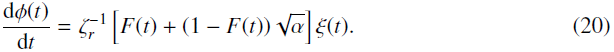

This model represents a first step towards investigating the mechanism of reorientation in *R. sphaeroides*.

### 2.6 Numerical implementation

To verify our analytic results and gain quantitative insight where such results are not possible, we perform numerical simulations of each of the self-propelled particle models numerically. We employ the Euler-Maruyama (EM) method [26], a discrete-time approximation to the underlying equations. The EM method requires that we specify a simulation time step, Δ*t*, which must be sufficiently small to ensure numerical stability. The MATLAB code used to implement these simulations is provided as Electronic Supplementary Material and allows the simulation of each model as a special case of the run-and-active-stop model (18)–(20). The run-only model (12)–(14) is simulated by setting *λ*_*s*_ = 0, and hence preventing stops. The run-and-stop model (15)–(17) is simulated by setting *α* = 1, in which case there is no additional reorientation force. All simulations carried out had a time step Δ*t* = 0.02 s and a total simulation time of 10 s. The other parameter values used were *T* = 300 K, *η* = 10^−3^ N sm^−2^ and *k* = 1.38 × 10^−23^ kg m^2^ s^−2^ K^−1^. We simulate 5000 tracks each time the algorithm is run. To ensure consistency, the results of the simulation are compared with the analytic expressions for the moments derived in the Electronic Supplementary Material.

### 2.7 Experimental methods

Full details of the experimental protocol used to generate the bacterial tracking datasets used in this study, as well as the algorithm used to extract cell tracks, are given in Rosser et al. [18]. Strains used were *R. sphaeroides* WS8N (wild type) [27], JPA 1353 (non-chemotactic) deleted for chemotaxis operons 1, 2 and 3, cheBRA, and CheY4 [6] and JPA 467 (non-motile) [28]. Cultures were grown aerobically in succinate medium to mid-log phase, at which point they were maximally motile [27].

The analysis method used here enables the non-parametric annotation of reorientation phases in the tracks and is compatible with any motile behaviour that is well-approximated by the run-and-stop model described previously [18]. The approach takes advantage of the availability of non-chemotactic and non-motile mutants to gain empirical knowledge of the appearance of running and stopping phases in the observed motion. The methods are based on a modification to the hidden Markov model, and are applicable to any bacterial species where such mutants exist and sufficiently long reorientation events are discernible using video microscopy. A Python implementation of the analysis software is freely available to download at http://www.2020science.net/software/bacterial-motility-analysis-tool. The tracking dataset is provided as Electronic Supplementary Material.

In this study, we used the same protocol described in the referenced article, with the exception of a minor modification to the censoring process. We removed the top 10% of tracks based on median curvature, compared with the 5% removed in the original study. This results in the removal of an additional 85 tracks from the non-chemotactic mutant dataset and 160 tracks from the wildtype dataset, leaving 1531 and 2878 tracks, respectively. The implications of jagged tracks in the dataset are discussed in [18]; in particular, their presence generates a large number of spurious reorientation events and large between-frame angle changes. Our modified censoring stage is necessary as the analyses in the original study are more robust to such effects than those in this study. We find that failure to filter these tracks results in very severe departures from linearity in the mean squared angle change (see Section 3.2). We discuss this in more detail in the Electronic Supplementary Material.

## 3 Results

In this section, we analyse each model to obtain experimentally-testable predictions. We make several independent comparisons between our theoretical results and both simulated and experimental data. We shall use various statistics relating to the experimental data, including the displacements observed between consecutive sample points, denoted framewise displacements (FDs), the angle change between consecutive pairs of sample points, denoted framewise angle changes (FACs), and the angle change observed over the course of a reorientation event, denoted stopwise angle changes (SACs).

As noted, the experimental data used for comparison are obtained from a tracking study in *R. sphaeroides* in which wildtype and mutant strains were observed. Before commencing analysis of the models, we first consider the simple case of the non-motile strain. This is an important consistency check, since the foregoing theory dictates that the observed motion of such cells will be fully described by Brownian translational diffusion. It also provides a well-known example with which to introduce some of the mathematical concepts used throughout this study to obtain summary statistics from SDE models.

### 3.1 Translational diffusion in non-motile bacteria

The translational motion of a non-motile bacterium undergoing isotropic translational diffusion in a fluid is given by equation (6). A common means of testing whether observed motion is diffusive involves the time evolution of the mean squared displacement (MSD). This quantity is derived in the Electronic Supplementary Material and is given by

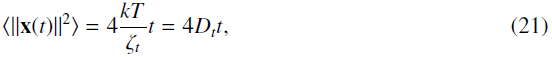

where *D*_*t*_ = *kT*/*ζ*_*t*_ is the translational diffusion coefficient. The MSD of the non-motile dataset is shown in Figure 4(a). The plot does not appear linear for *t* ≲ 0.2 s, so we fit to data outside this region in order to estimate *D*_*t*_. We attribute this departure from linear behaviour to artefacts generated by the tracking process: the tracking algorithm may fill in any missed detections by assuming that the intervening motion took place with a constant velocity, leading to an overestimation of the FDs. This would lead to a non-linear MSD with a steeper slope, as observed. This effect is only present over short time intervals, as the tracking algorithm may only fill in short gaps. At longer time intervals, linear behaviour is resumed.

We carry out an additional check that the observed motion is approximately diffusive by plotting the distribution of observed cell displacements *R*(*τ*) for various time intervals *τ*, achieved by successively downsampling the data. This distribution is derived in the Electronic Supplementary Material and has probability density function (PDF)

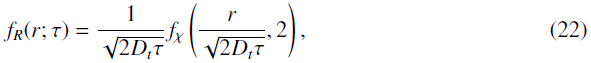

where *f*_*χ*_(·, *k*) denotes the PDF of the *χ* distribution with *k* degrees of freedom. We fit PDF (22) to the observed data to extract an estimate of *D*_*t*_. The observed and fitted distributions are shown in Figure 4(b) for three values of *τ*. The fit becomes progressively better as *τ* increases, although it shows good qualitative agreement for all time lags. The observed distributions further support our hypothesis that the departure from the theoretically predicted behaviour is due to an overestimation of step lengths by the tracking algorithm: the observed distributions show a heavier tail than predicted for all values of *τ*.

**Figure 4.**
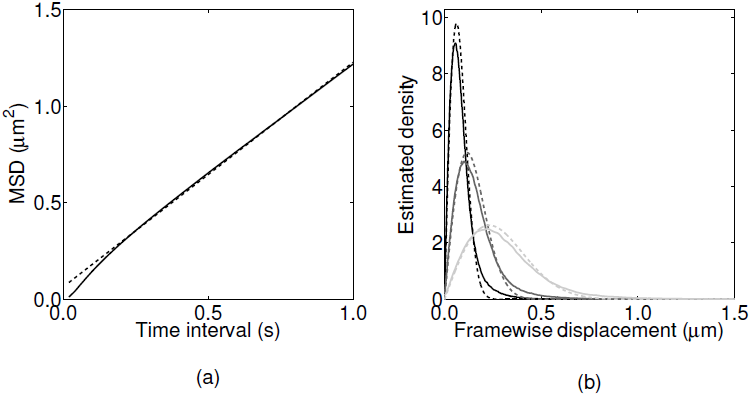
Extracting characteristics of noise from the *R. sphaeroides* non-motile dataset. (a) The observed MSD (solid line) with linear fit (dashed line). The slope of the fit is 1.16 *μ*m^2^ s^−1^. This Figure reproduces Figure S12 in Rosser et al. [18]. (b) The estimated PDF of framewise displacements (solid line), with overlaid best fit to the *χ* distribution (dashed line). The three shades denote different values for the sampling interval, *τ*. Black: *τ* = 0.02 s; dark grey: *τ* = 0.04 s; light grey: *τ* = 0.1 s.

Table 1 displays the estimates of *Dt* calculated using a linear fit to the MSD, and by fitting (22) to the observed distribution of displacements. Table 1 also shows the theoretical value of *Dt* calculated by approximating a bacterium as a sphere and applying equation 5. We take *r* = 1 *μ*m and the values for *k*, *T* and *η* given in Section 2.6, which yields *D*_*t*_ = 0.22 *μ*m^2^ s^−1^. The various estimates are in good agreement with the theoretical value, especially considering that the radius varies between cells in the population. For example, recalculating the theoretical value with *r* = 0.75 *μ*m, which is still a realistic value for a bacterium of slightly smaller size, we obtain *D*_*t*_ = 0.29 *μ*m^2^ s^−1^.

**Table 1.** Theoretical and estimated values of the translational diffusion coefficient of the non-motile strain of *R. sphaeroides* using data obtained by Rosser et al. [18].

| Method | $D_t$ ($\mu\text{m}^2 \text{s}^{-1}$) |
| --- | --- |
| Theoretical | 0.22 |
| MSD linear fit | 0.29 |
| $\chi$ fit, $\tau = 0.02 \text{ s}$ | 0.19 |
| $\chi$ fit, $\tau = 0.04 \text{ s}$ | 0.26 |
| $\chi$ fit, $\tau = 0.1 \text{ s}$ | 0.31 |

### 3b Run-only model

#### 3.2.1 Theoretical results

In order to validate our simulation, we begin by comparing numerical and analytic results for the run-only model. Figure S1(a) shows 20 sample trajectories obtained by simulating the run-only model (12)–(14) numerically, as described in Section 2.6. Analytic expressions for the mean and standard deviation of the *x* and *y* coordinates, which are derived in the Electronic Supplementary Material, are plotted in Figure S3(b). The results from a simulation are essentially indistinguishable and are thus not shown for brevity.

We now consider the angle change, *ϕ*(*t*), whose first and second moments are derived in the Electronic Supplementary Material. Our measured angle change is not equivalent to the true underlying change in orientation, as illustrated in Figure 5(a). Figure 5(b) shows the MSAC of an ensemble of simulated particles, computed using the two different methods. The solid line is calculated using the true underlying orientation, *ϕ*(*t*). This line agrees well with the analytic expression

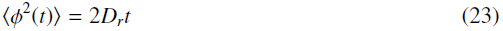

derived in the Electronic Supplementary Material. The dashed line in Figure 5(b) represents the *measured* MSAC, calculated by estimating angle changes from the recorded positions, which we denote by *ϕ*_meas_. The measured MSAC also scales linearly with time, but with a constant of proportionality that is 60% of that in the true MSAC. Misuse of the MSAC, by assuming that *ϕ*(*t*) ∼ *ϕ*_meas_(*t*) and using the latter quantity to compute *D*_r_, could therefore lead to a substantial underestimate of the rotational diffusion coefficient.

**Figure 5.**
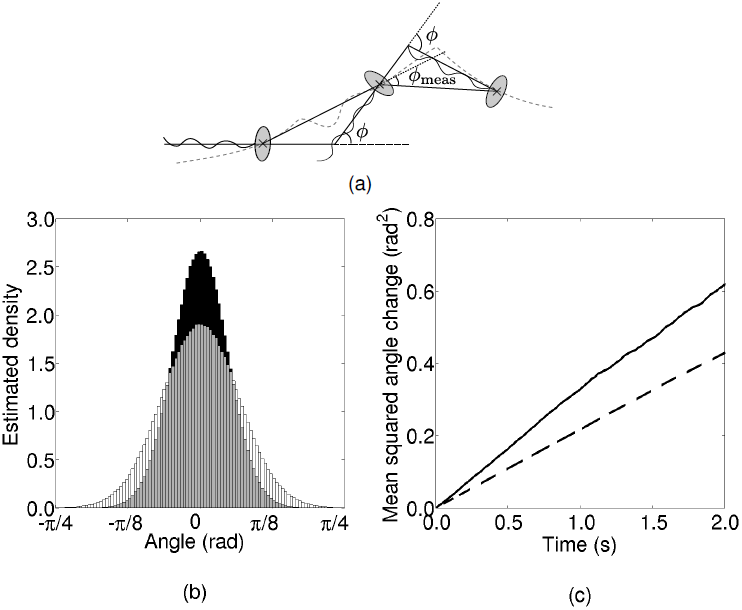
(a) Illustration of the difference between the true underlying angle change, *ϕ*, and the measured angle change, *ϕ*_meas_, of a swimming bacterium. The grey dashed line denotes the true trajectory of the bacterium. (b) Histograms of the measured angle changes, *ϕ*_meas_, of simulated particles in the run-only model for different values of the sampling time step *τ*. (Black) *τ* = 0.1 s, (white) *τ* = 0.2 s. (c) The observed MSAC over time for simulated tracks. (Solid line) angle change computed using the true underlying orientation of the simulated particles, (dashed line) angle change computed by calculating the framewise angle change from particle positions.

Just as for the MSD, it is possible to check not only the MSAC, but also the distribution of angle changes, against data. As given by (23), the true distribution of *ϕ*(*t*) is a wrapped normal, with variance proportional to time. It is not, however, obvious how *ϕ*_meas_(*t*) is distributed. To address this question, we simulate a particle undergoing a run-only process with rotational diffusion. Figure 5(c) shows the observed distribution of the measured angle changes for two different values of the sampling time step *τ*. We find that the distribution is indiscernible from the normal distribution in each case.

#### 3.2.2 The effect of cell geometry on rotational diffusion

We now consider how the geometry of the cell body affects the role of rotational diffusion on bacterial motility. We assume that there is little variation in the length of the equatorial (shorter) semi-principal axis in bacteria, and hence consider cells with a constant equatorial radius, *b* = 1 *μ*m. We then vary the length of the axial radius *a*, hence the axial ratio *ρ*, to study a range of prolate ellipsoids (for which *a* > *b*). Figure 6(a) shows the effect of varying *ρ* on the rotational drag coefficient, *ζ*_*r*_. The solid line corresponds to the drag coefficient for rotation in the equatorial axis, and the dashed line indicates the drag coefficient for the sphere of equivalent volume. This plot demonstrates that as the axial ratio increases, the rotational drag coefficients for a sphere and for an ellipsoid of equal volume diverge from each other, and ellipsoidal bacteria are more stable to rotational diffusion than spherical bacteria. Figure 6(b) shows the variation of the diffusion coefficient for rotation in the equatorial axis with *ρ* for an ellipsoid of fixed volume. We see that the rotational diffusion coefficient decreases with *ρ*, again indicating that an ellipsoidal geometry stabilises the cell towards Brownian reorientation.

**Figure 6.**
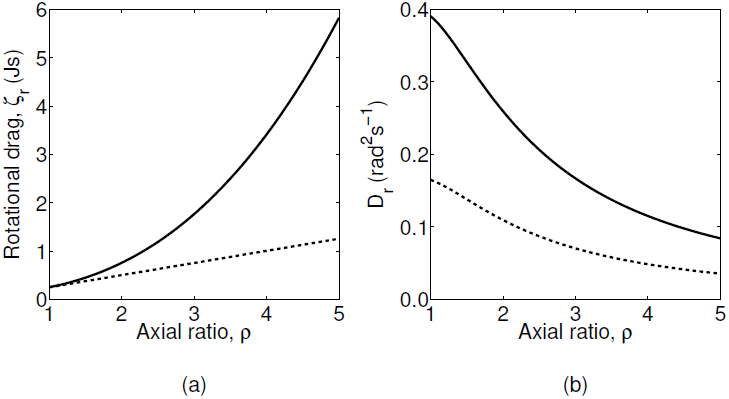
(a) The variation of the rotational drag coefficient *ζ* with *ρ* for a prolate ellipsoid with equatorial radius of length 1 *μ*m. The solid line represents the rotational drag coefficient for rotation in the equatorial semi-principal axis, while the dashed line shows the rotational drag coefficient for a sphere of equivalent volume. (b) The variation of the rotational diffusion coefficient for rotation in the equatorial semi-principal axis with *ρ* for a prolate ellipsoid of fixed volume. The radius of the equivalent sphere is 0.75 *μ*m in the case of the solid line, and 1 *μ*m in the case of the dashed line.

We further illustrate this point by returning to the work of Mitchell [13], who derived equations for the minimum useful swimming speed for bacteria in terms of the size of the bacterium. For a chemoattractant with molecular translation diffusion coefficient *D*_*m*_ ≈ 1000 *μ*m^2^ s^−1^, Mitchell defined a characteristic length, 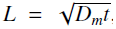, over which the change in the concentration of chemoattractant is sufficiently large to be detected by the bacterium in time *t*. The minimum useful speed is given by *v*_min_ = *L*/*t*. Hence, from (23), we have

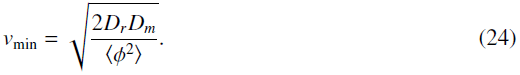

We interpret this minimum distance as the mean distance travelled before rotational diffusion causes an orientation change of *π*/2 [13]. We solve equation (24) with 〈*ϕ*^2^〉 = (*π*/2)^2^, indicating that a cell must travel a distance *L* before its mean reorientation angle exceeds *π*/2. We note that this interpretation is an approximation, as the equality 〈*ϕ*^2^〉 = (*π*/2)^2^ does not imply that 〈|*ϕ*|〉 = π/2. Figure 7(a) shows the variation of *v*_min_ with axial ratio, *ρ*, for cells with various equatorial radii, *b*. Most wildtype *R. sphaeroides* cells have an equatorial radius in the range 0.5 *μ*m < *b* ≲ 1 *μ*m [25, 29]. Figure 7(b) shows the mean swimming speed for each track in the *R. sphaeroides* non-chemotactic bulk dataset. The absence of measurements of the diameters of tracked cells limits any quantitative analysis of these data, however we are able to determine that the plots in Figure 7 are consistent in the sense that most cells swim sufficiently quickly to meet the minimum useful speed, assuming that 0.5 *μ*m ≲ *b* ≲ 1 *μ*m and *ρ* ≳ 1.5.

**Figure 7.**
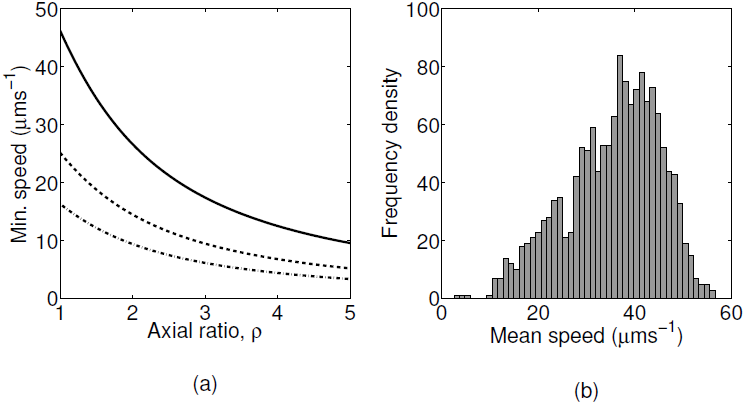
(a) The variation of minimum useful speed, *v*_min_, with axial ratio for ellipsoidal bacteria with different fixed equatorial radii, *b* = 0.5 *μ*m (solid line), *b* = 0.75 *μ*m (dashed line), and *b* = 1 *μ*m (dotted line). (b) Histogram of the mean speeds for each track in the *R. sphaeroides* non-chemotactic bulk dataset.

#### 3.2.3 Comparison with experimental data

We now seek to estimate the rotational diffusion coefficient, *D*_*r*_, in the run-only model for the non-chemotactic strain of *R. sphaeroides*. These tracks are approximately described by our run-only model. The MSAC in this model grows linearly with time, as given by (23). Furthermore, we expect that angle changes should follow a wrapped normal distribution with zero mean and variance equal to the MSAC. Figure 8 shows the comparison of the non-chemotactic dataset with the predictions of the model. As for the experimentally observed MSD, the MSAC is ap-proximately linear for *τ* ≳ 0.2 s. Estimating *D*_*r*_ from the linear portion of the data in Figure 8(a) yields *D*_*r*_ = 0.13 rad^2^ s^−1^. However, as discussed in Section 3.2.1, the use of FACs as an approximation to the true orientation *ϕ* is expected to underestimate *D*_*r*_ by around 60%. Correcting for this error, we obtain an estimate of *D*_*r*_ = 0.21 rad^2^ s^−1^.

**Figure 8.**
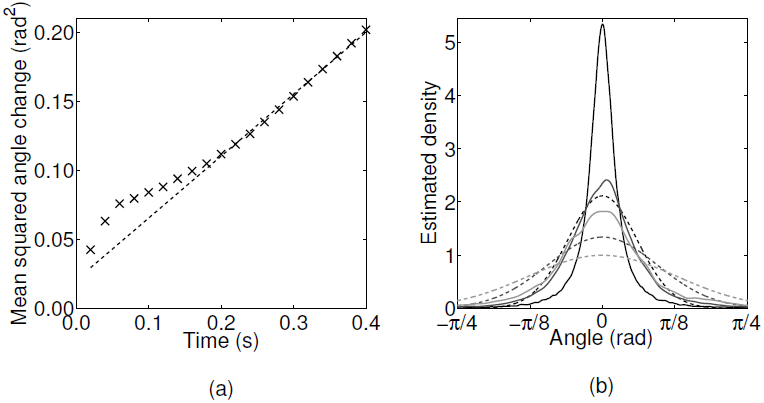
Extracting characteristics of noise from the *R. sphaeroides* non-chemotactic dataset obtained by Rosser et al. [18]. (a) The observed MSAC (black crosses) with linear fit (dashed line). The slope of the fit is 0.255 rad^2^ s^−1^. (b) The estimated PDF of angle changes (solid line), overlaid with the wrapped normal distribution with variance computed from the data (dashed line). The three shades denote different values for the sampling interval, *τ*. Black: *τ* = 0.02 s; dark grey: *τ* = 0.1 s; light grey: *τ* = 0.2 s.

Hence we may estimate the rotational diffusion coefficient from the data in Figure 8(a), giving *D*_*r*_ = 0.13 rad^2^ s^−1^.

Figure 8(b) shows the observed distribution of angle changes for three values of the sampling time step, *τ*, overlaid with the predicted wrapped normal distribution with zero mean and variance calculated from the data. There is a significant discrepancy between the observed and predicted distributions, for which we offer two possible explanations. First, errors in finding cell centroids, in addition to artefacts generated by the tracking algorithm, may contribute to the departure of the results from the predicted normal distribution. In particular, the tracker developed by Wood et al. [30] that is used to generate the experimental datasets in [18] is based on a constant velocity motion model with additional Gaussian noise terms. This is expected to over-estimate small observed FACs, leading to a more peaked distribution, as observed. Second, our computation of *ϕ*(*t*) is based on the assumption that FACs may be used to approximate changes in the orientation of the cell. Whilst this cannot explain the non-normal distribution of observed FACs, it is expected to produce a distribution with lower variance than the true distribution (see Figure 5(b)), which is similar to that seen in Figure 8(b).

The theoretical value of the rotational diffusion coefficient is calculated using equation 9. Approximating a bacterium as a sphere of radius *r* = 1 *μ*m, we obtain *D*_*r*_ = 0.16 rad^2^ s^−1^, in close agreement with the value estimated from the data. We note, however, that the theoretical estimate is highly sensitive to the value of *r*; for example, choosing *r* = 0.75 *μ*m gives *D*_*r*_ = 0.39 rad^2^ s^−1^.

### 3.3 Run-and-stop model

#### 3.3.1 Theoretical results

As for the run-only model, we begin by comparing numerical and analytic results. Analytic expressions for the first and second moments of the angle change *ϕ*(*t*) and *x* and *y* coordinates satisfying the run-and-stop model (15)–(17) are derived in the Electronic Supplementary Material. Figure S2 shows the analogous plots to those in Figure S1 for the run-and-stop model. The mean and standard deviation of *x*(*t*), and the standard deviation of *y*(*t*), have lower magnitudes than the run-only analogues as stopping phases cause the cell to halt sporadically, which reduces the extent of travel and dispersion.

#### 3.3.2 Comparison with experimental data

We now compare the predictions of the run-and-stop model with experimental data, in order to assess how well the model captures the observed motile behaviour of *R. sphaeroides*. For this purpose, we use the wildtype dataset. We focus on the predicted variation of the variance of SACs with stop duration. According to the run-and-stop model, the evolution of the orientation angle *ϕ* during both running and stopping phases is described by equation (17), which states that angle changes follow the wrapped normal distribution with zero mean and variance equal to 2*D*_*r*_*τ*, where *τ* is the stop duration. The theoretical distribution of SACs is shown in Figure 9(a) for several different stop durations.

**Figure 9.**
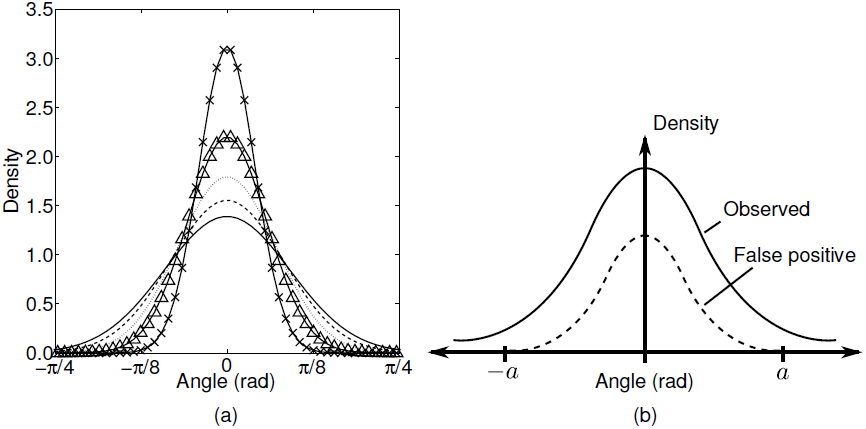
(a) The theoretical distribution of stopwise angle changes, as predicted by the run-and-stop model, for stops of 0.1 s (×), 0.2 s (Δ), 0.3 s (dotted line), 0.4 s (dashed line), and 0.5 s (solid line). (b) An illustration of the problem of false positives due to analysis artefacts in the distribution of stopwise angle changes. The exact form of the dashed line distribution is unknown, but we can estimate the cutoff value, *a*, reasonably well.

A direct comparison of Figure 9(a) with the experimental data requires that we estimate the variance of the observed SACs, denoted 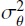, for a variety of stop durations. This process is complicated by the presence of artefacts from the analysis procedure, as discussed in Rosser et al. [18]. The problem is summarised in schematic form in Figure 9(b). The observed density of SACs (solid line) has a substantial false positive component (dashed) line, which will skew our estimate of the variance. This is quantified in the simulation study by Rosser et al. [18, Figure 5], who show that the density of false positives decreases with increasing stopwise angle change. Estimation of the exact distribution of false positives in the experimental dataset is not possible, so we instead define an *acceptance region*, |*θ*| > *a*, in which we assume that the density of false positives is negligible. Based on the results in [18], we choose *a* = *π*/2. We first bin the observed stops by their duration. For each group, we estimate the total density of SACs in the acceptance region in two ways:

1. Assume no false positives, by taking the number of angle changes in the acceptance region divided by the total number of angle changes.
2. Use the simulated dataset from Rosser et al. [18] to estimate the false positive level outside the acceptance region, and hence to estimate the density within the acceptance region, taking false positives into account.

Method 1 provides a lower bound on the density of SACs in the acceptance region, and therefore on the value of 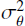, since the presence of false positives outside the acceptance region is ignored. These false positives are clustered around the mean (i.e. zero), hence they artificially reduce the apparent variance. Method 2 constitutes an improved estimate that is corrected in an approximate fashion for false positives; note that this is not an upper bound, nor is it necessarily an accurate estimate, however it is likely to be more accurate than method 1. The estimated proportion of false positives outside the acceptance region in the simulated dataset is shown in Table 2. We use the data simulated with the level of noise given by *D* = 0.288 *μ*m^2^ s^−1^, as this value is in close agreement with the estimates of *D*_*t*_ listed in Table 1.

**Table 2.**
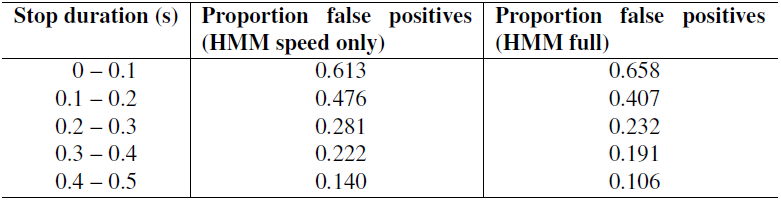
The proportion of false positives outside the acceptance region for the simulated dataset generated by Rosser et al. [18].

| Stop duration (s) | Proportion false positives<br>(HMM speed only) | Proportion false positives<br>(HMM full) |
| --- | --- | --- |
| 0 – 0.1 | 0.613 | 0.658 |
| 0.1 – 0.2 | 0.476 | 0.407 |
| 0.2 – 0.3 | 0.281 | 0.232 |
| 0.3 – 0.4 | 0.222 | 0.191 |
| 0.4 – 0.5 | 0.140 | 0.106 |

We now seek an expression linking the total density in the acceptance region to the variance of the wrapped normal distribution with zero mean. The PDF of this distribution is given by

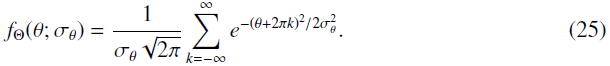

Note that this distribution is symmetric about *θ* = 0. The total density in the acceptance region is thus given by

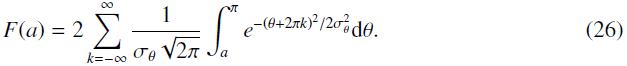

Each term in the summation is given by

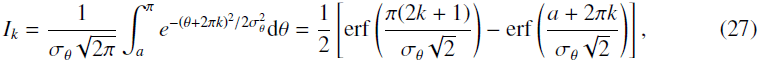

where erf(·) denotes the error function. Substituting (27) into (26), we obtain

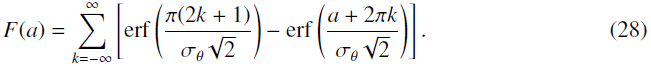

Having obtained an estimate for *F*(*a*) using either of the methods described above, we use trust-region constrained numerical optimisation [31] to find *σ*_*θ*_ from (28). The results of this calculation, along with the theoretically predicted result for an ellipsoidal bacterium with an equatorial diameter of 0.5*μ*m, are shown in Figure 10. The geometry of the model bacterium is selected as an approximate match to the true dimensions of *R. sphaeroides*. The discrepancy between the lower bound estimate of 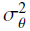 and the theoretical value is striking; the theoretical variance is an order of magnitude smaller than the lower bound. This demonstrates that *R. sphaeroides* does not reorientate passively by rotational Brownian diffusion: the bacterium is too large for this to be an effective mechanism for reorientation.

**Figure 10.**
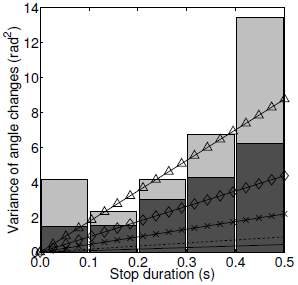
The lower bound (black bars) and revised estimates (grey bars) of the variance of stopwise angles, classified by stop duration, in the wildtype *R. sphaeroides* bulk dataset. Overlaid lines show theoretical results for the variance of stopwise angle changes predicted by the run-and-active-stop model for an ellipsoidal bacterium, with *b* = 0.5 *μ*m, *ρ* = 2, and *α* = 1 (solid line, equivalent to the run-and-stop model with no active reorientation force), *α* = 2 (dashed line), *α* = 5 (×), *α* = 10 (◊), and *α* = 20 (Δ).

### 3.4 Run-and-active-stop model

Figure 10 also shows predictions from the run-and-active-stop model. The multiplicative factor α increases the variance of angle changes linearly. The data are too noisy to permit us to fit α accurately, but the data suggest that wildtype *R. sphaeroides* reorientate between 5 and 20 times more rapidly than predicted by the run-and-stop model. This result strongly suggests that there is active movement, presumably slow or transient motor rotation, during a stop, which increases the rate of reorientation.

## 4 Discussion

In this study, we have considered the role of Brownian diffusion in flagellar-mediated bacterial motility. We focused on rotational diffusion, ignoring translational diffusion, since it is the former that leads to fundamental limitations in the way that bacteria propel themselves through a liquid medium. The absolute magnitude of the perturbation from a straight line trajectory caused by rotational diffusion varies with the speed at which a cell travels, whereas translational diffusion has a fixed effect regardless of the rate of propulsion. This work is motivated by the need to reconcile the idealised VJ model, considered in Rosser et al. [32], with the immediately apparent departure from the model in experimentally observed tracks (see Figure 1) and the differences seen in experiments with tethered cells and free-swimming cells.

We used an overdamped Langevin equation, in the context of a self-propelled particle model, to model rotational diffusion. This approach was motivated by a related study by ten Hagen et al. [19]. We described three minimal models for bacterial motility in a planar domain. This geometrical simplification was justified by our use of experimental data of free-swimming tracks within a focal plane. The three models were chosen to describe the motion of *R. sphaeroides* wildtype and non-chemotactic mutant bacteria.

We tested several hypotheses arising from our models against the experimental data, comparing the observed PDF of angle changes with the theoretically predicted distribution, demonstrating that the mean swimming speeds of *R. sphaeroides* agree well with theoretical minimum bounds, and testing a model for the poorly-understood mechanism of reorientation in this monoflagellate. All experimental data were analysed using the methods presented in Rosser et al. [18].

While similar studies have been undertaken in other biological systems, such as human dermal keratinocytes [33] and the amoeba *Dictyostelium discoideum* [34], to our knowledge our work is the first such comparative study in bacteria. Furthermore, in [34], the authors take a more generalised approach to their analysis, allowing departures from the standard Langevin formulation. It is not clear how this more general approach relates to the underlying physical theory of motion, or the biological processes involved. In contrast, in the present study we have robustly tested our model predictions.

Our results show good agreement between the theoretical and observed translational diffusion coefficients *D*_*t*_, inferred using non-motile strain data. In particular, we found that the *distribution* of displacements match the theoretical predictions well. This test is often ignored in favour of simply testing the MSD for linearity [35, 36]. When data are scarce, this is an acceptable compromise, however our analysis shows that a full comparison of the distribution, when possible, is a useful test.

We found that the observed dimensions and shape of *R. sphaeroides* in the experimental data are consistent with the mean speeds observed in the tracks. We also found good agreement between the non-chemotactic dataset and the predictions of the run-only model, quantified through comparison of the MSAC. The rotational diffusion coefficient *D*_*r*_ estimated in this way agreed with the theoretically predicted value. However, we identified significant departures from the predicted distribution of angle changes, which we attributed to errors in the image analysis and tracking procedures, in addition to the method used to estimate FACs. Regarding this latter point, we demonstrated the difference between the true orientation of a bacterium, which is not observable using present experimental protocols, and an approximation that is easily computed from the tracking data. Assuming that these two quantities are equal leads to an underestimate for *D*_*r*_.

In order to compare the predictions of the two models incorporating stochastic stopping events with our wildtype data, we needed to circumnavigate the issue of artefacts in the dataset. The simulation study carried out in Rosser et al. [18] was of great utility in this regard. Guided by this, we were able to demonstrate the important biological result that *R. sphaeroides* cannot reorientate purely passively by rotational diffusion. This was clear from the strong incompatibility between the predictions of the run-and-stop model and the observed data. An important contribution of our work is in providing strong novel evidence to reinforce this conclusion. To the authors’ knowledge, this study is the first to contain robust analysis of tracking data with reference to a well-stated model to address the question of the mechanism of reorientation in bacteria.

Motivated by this discrepancy, we also compared the experimental data with the predictions of the run-and-active-stop model, in which a stochastic rotational force leads to an increased rate of reorientation during stops. Significant variability in the data, coupled with a need to compensate for potential bias introduced by the analysis process, prevents us from estimating the strength parameter α with a high degree of confidence in the present study.

We do not consider the precise nature of the active rotational force in the present work, as it is not the goal of our analysis, however several previous studies suggest plausible mechanisms. The mechanism of reorientation in *R. sphaeroides* and related bacterial species has been discussed before in the literature, with no strong consensus [6, 7, 13, 16]. The hypothesis that *R. sphaeroides* exhibits some form of active reorientation mechanism was briefly discussed by Armitage et al. [7]. The authors present a study in which they observed the various flagellar conformations of swimming and stopping *R. sphaeroides* cells. In particular, a conformation was observed in stopped cells in which “once coiled against the cell body, the flagellum often slowly rotated”. In the context of the present work, we speculate that such a low-frequency movement of the flagellum during a stopping phase may lead to enhanced rotational diffusion. A further speculative mechanism involves the elastic relaxation of the flagellum or the hook complex linking it to the motor upon entering a stop phase.

A recent study has shown that the number of engaged motor stators increases with the force applied to the filament. It seems probable that a certain rotational force is required to drive reformation of the functional flagellum following a reorientation phase, causing a period of rotation of the relaxed filament before that torque is reached. Previous work on the mechanism of reorientation in *R. sphaeroides* [6] suggests that the braking torque during a stop is greater than the stall torque of a disengaged motor, but used a tethered cell assay with filaments under increasing loads, and therefore may have artificially increased the number of engaged stators.

Further detailed microscopy studies of individual free-swimming cells and their flagella are required to elucidate the mechanisms involved, as much detailed information on flagellar dynamics is gained from this type of study [37, 38].

We did not include the additional frictional effects of the flagellum in the present work, which may be expected to reduce the reorientation rate even further [13]. This additional flagellum drag is expected to be less significant in *R. sphaeroides* than *E. coli*, as the former posses a single, thin flagellum, compared with the flagellar bundle in peritrichous bacteria such as *E. coli*. Furthermore, in order to reduce the number of free variables in this study, we computed theoretical results of the run-and-stop and run-and-active-stop models using a single representative bacterial cell geometry. Longer, thinner ellipsoidal bacteria would experience reduced rotational diffusion.

By demonstrating the utility of minimal models and quantitative comparison with the largest available tracking dataset of free-swimming *R. sphaeroides* to address key biological questions, the present work represents an important contribution to the field of bacterial motility. There are many opportunities for further investigation. We identified several departures from the theoretical predictions, particularly when comparing the non-chemotactic dataset with the run-only model. We suggested that these differences may be explained by artefacts introduced by image analysis and cell tracking, or due to the method of calculating the angle change. Further studies could test whether these departures contain useful information about the biological processes underlying bacterial motility, or are indeed due to artefacts introduced in the various analysis stages.

A related consideration is variation in swimming speeds across individuals. Several studies have noted the highly heterogeneous nature of bacterial populations in terms of swimming speeds [7, 39, 40]. In our models, we fixed the swimming speed; an obvious extension is to replace this with a fluctuating quantity. Random velocities have been considered in the context of random walks [41], but have not been coupled with a model of rotational diffusion. This may further explain the departure of the experimental results from the run-only model predictions. It is, however, unclear how the random velocities should be distributed: a *population*-level distribution of velocities was measured by Rosser et al. [18], but a distribution on an *individual-level* would be required for this extension.

## Acknowledgments

The authors thank Prof. Philip K. Maini for valuable feedback on this work. G.R. is supported by an EPSRC-funded Life Sciences Interface Doctoral Training Centre Studentship (EP/ES0160S/1). G.R. and A.G.F. are funded by EPSRC (EP/I017909/1) and Microsoft Research Cambridge.

